# Associations between recurrent mutations and blast immunophenotype in acute myeloid leukemia

**DOI:** 10.1101/2021.04.14.439800

**Authors:** Kateřina Kuželová, Barbora Brodská, Jana Marková, Martina Petráčková, Šárka Ransdorfová, Zdenka Gašová, Cyril Šálek

## Abstract

The immune system undoubtly plays an important role in final elimination of residual leukemic cells during acute myeloid leukemia (AML) therapy. However, the anti-leukemia immune response can be inhibited by a variety of mechanisms enabling immune escape and eventual disease relapse. We analyzed selected markers of immune escape on AML cells at diagnosis (N = 53) and used them for hierarchical clustering analysis, which yielded distinct clusters with different incidence of mutations in nucleophosmin 1 (NPM1) and in the methyltransferase DNMT3A. More detailed analysis showed that in the absence of DNMT3A mutation, NPM1 mutation is associated with decreased HLA expression and also with low levels of other markers (CLIP, PD-L1, TIM-3). On the other hand, samples with concomitant DNMT3A mutation had high CLIP surface amount suggesting reduced antigen presentation. Higher CLIP exposition was also found in patients with internal tandem duplications in FLT3 (FLT3-ITD). TIM-3 transcript correlated not only with TIM-3 protein surface amount, but also with CLIP and PD-L1, suggesting acquisition of a complex immunoresistant phenotype. Our results indicate that AML genotype is to some extent related to the blast immunophenotype, and the established predictive values of particular mutations might also reflect an inherent cell resistance to the immune system.

## Introduction

Immunotherapy is an attractive therapeutic approach, which could complement the current treatment strategies in acute myeloid leukemia (AML). The most important challenge in AML treatment is associated with frequent disease relapses after chemotherapy. The curative potential of allogeneic bone marrow transplantation indicates that a functional immune system is able to eradicate residual leukemia cells. However, the anti-tumor immune response is often suppressed by means of different mechanisms, which are currently intensively investigated, especially in solid tumors. Evidence of immune system impairment is also available for AML [1–6]. Immunotherapeutic modalities include among others blocking antibodies against inhibitory receptors, bispecific antibodies, adoptive T-cell or natural killer (NK) cell therapy, or administration of T-cells with chimeric antigen receptors (CAR T-cells) [6–8].

A number of mechanisms could be involved in AML blast resistance to the immune system. The inhibitory receptors PD-L1 and PD-L2 bind to PD-1 on cytotoxic T-lymphocytes and prevent the target cell lysis. Blocking antibodies against these so-called checkpoint inhibitors are in clinical trials for many tumor types, including AML, usually in combination with another treatment. Another mechanism of immune evasion consists in reduced HLA expression, which impairs antigen-specific recognition of tumor cells by CD4+ or CD8+ T-cells. The invariant chain peptide CLIP, which can be present on HLA molecules instead of antigens, indicates defective antigen presentation. Although CLIP essentially forms part of HLA class II, it can also be cross-presented on HLA class I [9]. In AML, CLIP down-modulation enhanced the immunogenicity of myeloid leukemic blasts and resulted in increased CD4+ T-cell responses [10]. CLIP presentation on the surface of leukemia blasts was related to worse outcome in AML [11]. TIM-3, which was originally uncovered as a marker of exhausted T-cells [12], is often present on AML blasts [13]. Its function in leukemia cells is only partially understood and is probably complex. TIM-3 enhances leukemia cell proliferation and resistance to apoptosis through autocrine or paracrine signaling [14]. It is also secreted in complex with galectin-9, which inhibits the activity of T-cells and thereby contributes to immune escape [15]. CD47 serves as a “don’t-eat-me” signal to macrophages and its presence on AML blasts was associated with worse outcome [16]. CD47 targeting was suggested as a possible therapeutic anti-cancer approach [17,18].

Activation of signaling pathways regulating the immune escape might form an integral part of oncogenic transformation. We and others have previously revealed some associations between recurrent mutations found in AML and specific immunophenotypic features of leukemia cells. For example, nucleophosmin 1 (NPM1) mutation leads to neoantigen formation as well as to aberrant cytoplasmic localization of the protein product, possibly leading to more efficient NPM1 processing and presentation on HLA molecules. Several HLA class I alleles were underrepresented in patients with NPM1 mutation [19,20], possibly due to spontaneous cure of individuals with HLA types suitable for presentation of NPM1-derived immunopeptides. Such peptides were indeed found in the immunopeptidome of AML blasts [21,22], and T-cells reactive against NPM1 were repeatedly detected [23–26]. NPM1 mutation was also associated with lower HLA-DR expression [27]. Among NPM1-mutated AML, specific protein expression pattern including HLA-DR positivity predicted worse outcome [28,29]. Furthermore, NPM1 was described to interact with PD-L1 promotor and to regulate PD-L1 gene transcription in a breast cancer model [30]. AML-associated NPM1 mutation results in decreased NPM1 amount in the cell nucleus and could prevent NPM1 from its regulatory roles in gene transcription. Lower PD-L1 amount thus could be expected in cases with NPM1 mutation. On the other hand, high PD-L1 expression, which was assessed on the transcript level, predicted worse overall survival specifically in patients with internal tandem duplications in Fms-like tyrosine kinase 3 (FLT3-ITD) [31]. FLT3-ITD was also associated with higher CD47 expression [16] and with higher positivity of CD4+ T-cells for TIM-3, as a marker of lymphocyte exhaustion [32].

The DNA methyltransferase 3A (DNMT3A) regulates gene transcription by promotor methylation. Loss-of-function mutations occurring in AML might enhance the transcription rate of genes, which are involved in the immune escape. DNMT3A mutation was actually associated with higher HLA-DR positivity [33] and with decreased methylation of PD-L1 promotor [34]. Furthermore, reduced DNMT3A expression, or inhibition of DNMT3A activity by decitabine correlated with higher PD-L1 expression in breast cancer [35], resp. in melanoma [36].

In this work, we analyzed selected indicators of immune evasion on AML blasts. Primary cells from peripheral blood were tested at the time of AML diagnosis for the presence of selected surface markers, which could be relevant for AML. Hierarchical clustering analysis yielded two distinct clusters with different incidence of NPM1 and/or DNMT3A mutations. We thus focused to possible impact of recurrent mutations on AML blast immunophenotype.

## Results

The basic characteristics of AML patients included in this study is given in Table 1. Mononuclear cell samples from the peripheral blood of AML patients with hyperleukocytosis were isolated from leukapheresis products obtained at diagnosis, prior to treatment. The amounts of HLA class I, HLA-DR, CLIP, PD-L1, PD-L2, TIM-3, and CD47 on the surface of leukemia blasts were measured by flow cytometry. PD-L2 was not detected in any of the first 20 samples and it was subsequently removed from the panel. As it is described in the Methods section, HLA class I and CD47 were quantified using the mean fluorescence intensity (MFI), whereas the other parameters are given as the fraction of positive cells (%). Flow cytometry results were well reproducible when the analysis was repeated from defrozen sample aliquots, except for TIM-3. Due to a low stability of this marker, we assessed TIM-3 also at the mRNA level.

**Table 1:** AML patient cohort characteristics. The karyotype and the mutation status of nucleophosmin 1 (NPM1), Fms-like tyrosine kinase 3 (FLT3), and DNA methyltransferase 3A (DNMT3A) were determined as a part of routine clinical procedures. NPM1 mutation was also checked by immunofluorescence. In one case, no mutation was detected by sequencing NPM1 exons 11+12, but the protein was localized in the cell cytoplasm (NPMc+). The samples denoted as “0” had no detected mutation in NPM1 exons 11+12 and the protein was localized in the nucleus/nucleoli. The presence of an internal tandem duplication in FLT3 (FLT3-ITD) or of DNMT3A mutation is indicated by “1” in the corresponding column. In patients with FLT3-ITD, the ratio of mutated and wild-type allele is specified. NA = not available.

| ID | Sex | Age at Dg | Karyotype | NPM1 mutation | FLT3-ITD (allelic ratio) | DNMT3A mutation |
| --- | --- | --- | --- | --- | --- | --- |
| 1 | F | 51 | 46,XX[6] | A | 1 (0.84) | 1 |
| 2 | M | 60 | NA | A | 1 (0.61) | 0 |
| 3 | M | 62 | 47,XY,der(3)t(3;17)(q29;q22),+8[2]/46,XY[6] | A | 0 | 1 |
| 4 | M | 68 | 46,XY,t(2;4)(p21;q28)[6] | 0 | 1 (8.01) | 1 |
| 5 | M | 60 | NA | A | 0 | 0 |
| 6 | F | 24 | 46,XX[22] | A | 1 (0.46) | 0 |
| 7 | F | 57 | 46,XX,t(2;11)(q21;p15)[7] | 0 | 1 (0.63) | 0 |
| 8 | M | 57 | 46,XY[20] | A | 0 | 0 |
| 9 | F | 34 | 46,XX[15] | Nm | 0 | 1 |
| 10 | M | 38 | 46,XY[9] | A | 0 | 0 |
| 11 | F | 58 | NA | 0 | 0 | 0 |
| 12 | F | 35 | NA | 0 | 0 | 0 |
| 13 | F | 48 | NA | A | 0 | 1 |
| 14 | F | 63 | 46,XX[22] | 0 | 1 (11.36) | 0 |
| 15 | M | 61 | 46,XY[22] | A | 1 (0.65) | 1 |
| 16 | F | 43 | 46,XX[22] | A | 1 (1.3) | 1 |
| 17 | M | 69 | 45,XY,-7[21]/46,XY[1] | 0 | 0 | 0 |
| 18 | M | 62 | 48,XY,+5,+7[11]/<br>47,XY,+4,del(9)(q11)[7]/46,XY[6] | 0 | 0 | 0 |
| 19 | M | 62 | NA | 0 | 0 | 1 |
| 20 | M | 65 | 46,XY[22] | A | 0 | 1 |
| 21 | F | 47 | 45,XX,-7,t(9;22)(q34;q11.2)[18]/<br>46,XX[4] | 0 | 0 | 0 |
| 22 | M | 22 | 46,XY[10] | A | 1 (0.43) | 1 |
| 23 | F | 23 | NA | A | 1 (0.91) | 0 |
| 24 | F | 42 | 46,XX[15] | NPMc+ | 0 | 0 |
| 25 | F | 31 | 46,XX,t(9;11)(p22;q23)[22] | 0 | 0 | 0 |
| 26 | F | 61 | 46,XX[10] | A | 1 (0.52) | 1 |
| 27 | M | 42 | 46,XY,t(5;11)(q35;p15)[16] | 0 | 1 (1.61) | 0 |
| 28 | F | 38 | NA | 0 | 0 | 0 |
| 29 | M | 41 | 46,XY[20] | 0 | 0 | 0 |
| 30 | F | 57 | 46,XX[20] | A | 0 | 1 |
| 31 | M | 45 | 46,XY[20] | 0 | 1 (0.62) | 1 |
| 32 | F | 38 | 46,XX,t(9;11)(p21;q23)[7]/<br>45,X,-X,t(9;11)(p21;q23)[7]/ | 0 | 0 | 0 |
|  |  |  | 47,XX,+8,t(9;11)(p21;q23)[6] |  |  |  |
| 33 | M | 57 | 46,XY[20] | A | 0 | 0 |
| 34 | M | 40 | 46,XY,t(6;11)(q27;q23)[6]/<br>47,XY,t(6;11)(q27;q23),+der(6)t(6;11)(q<br>27;q23)[5] | 0 | 0 | 0 |
| 35 | M | 56 | 46,XY[22] | A | 0 | 1 |
| 36 | M | 35 | 46,XY[21] | 0 | 1 (4.55) | 0 |
| 37 | F | 45 | 46,XX[20] | A | 1 (0.95) | 1 |
| 38 | M | 40 | 46,XY[20] | 0 | 0 | 0 |
| 39 | M | 42 | 46,XY[20] | 0 | 0 | 0 |
| 40 | F | 28 | 46,XX[12] | B | 1 (0.66) | 0 |
| 41 | M | 52 | 46,XY,t(X;13)(q27;q21)[24]/46,XY[1] | A | 0 | 0 |
| 42 | F | 47 | 46,XX[20] | A | 0 | 1 |
| 43 | F | 43 | 46,XX[23] | A | 0 | 0 |
| 44 | F | 21 | 46,XX[20] | A | 1 (10.4) | 1 |
| 45 | F | 65 | 46,XX,t(5;11)(p13.2;p13)[8]/46,XX[2] | A | 1 (32.5) | 1 |
| 46 | M | 62 | 46,XY[12] | ZQ | 1 (0.82) | 0 |
| 47 | F | 45 | 46,XX[15] | 0 | 1 (2.55) | 0 |
| 48 | F | 64 | 46,XX,t(5;11)(q35;p15)[22] | 0 | 1 (0.75) | 0 |
| 49 | F | 65 | 46,XXdel(5)(q14q35),der(7)t(7;21)(q22;<br>q?),t(10;17)(q?26;q?22),der(12)t(12;21)<br>(p?13;q?),der(15)t(1;15)(?;p?),del(21)(q<br>21;q22)[12]/46,XXdel(5)(q14q35),der(7)<br>t(7;21)(q22;q?),t(10;17)(q?26;q?22),der<br>(12)t(12;21)(p?13;q?),der(15)t(1;15)(?;p<br>?),-21[12]/47,XX,del(5)(q14q35),der(7)t<br>(7;21)(q22;q?),t(10;17)(q?26;q?22),der(1<br>2)t(12;21)(p?13;q?),der(15)t(1;15)(?;p?)<br>,+del(21)(q21;q22)[2] | 0 | 0 | 0 |
| 50 |  |  | 46,XY,inv(3)(p24q26),der(7)del(7)(p11.2<br>p22)inv(p11.2q11.23)del(q32q36)[20]/4<br>5,XY,inv(3)(p24q26),-7[5] | 0 | 0 | 0 |
| 51 |  |  | NA | 0 | 0 | 1 |
| 52 |  |  | 46,XY,t(5;11)(q35;p15)[25] | 0 | 1 (0.40) | 0 |
| 53 |  |  | 49,XY,+8,+8,+8[2]/46,XY[15] | A | 1 (0.04) | 1 |

### Hierarchical clustering analysis

Hierarchical clustering (www.wessa.net) of all the tested AML samples (N = 53) was performed using HLA class I, HLA-DR, CLIP, PD-L1, and TIM-3 values obtained from flow cytometry measurements. One group of closely similar samples (depicted as cluster 1 in Fig. 1A) was characteristic by low amounts of all the input parameters (Fig. 1B). On the opposite side, the cluster 2 contained samples with high expression of HLA class I, HLA-DR, and CLIP (Fig. 1B). Although TIM-3 mRNA levels were not used for cluster analysis, they were clearly distinct in the cluster 1 (usually low levels) compared to the other two groups (Fig. 1B). The majority of the remaining samples belonged to an intermediate group, except for one sample (ID 18), which had high levels of TIM-3 and of HLA class I. This patient had an altered karyotype with duplicated chromosomes 5 and 7 (Table 1).

**Figure 1:**
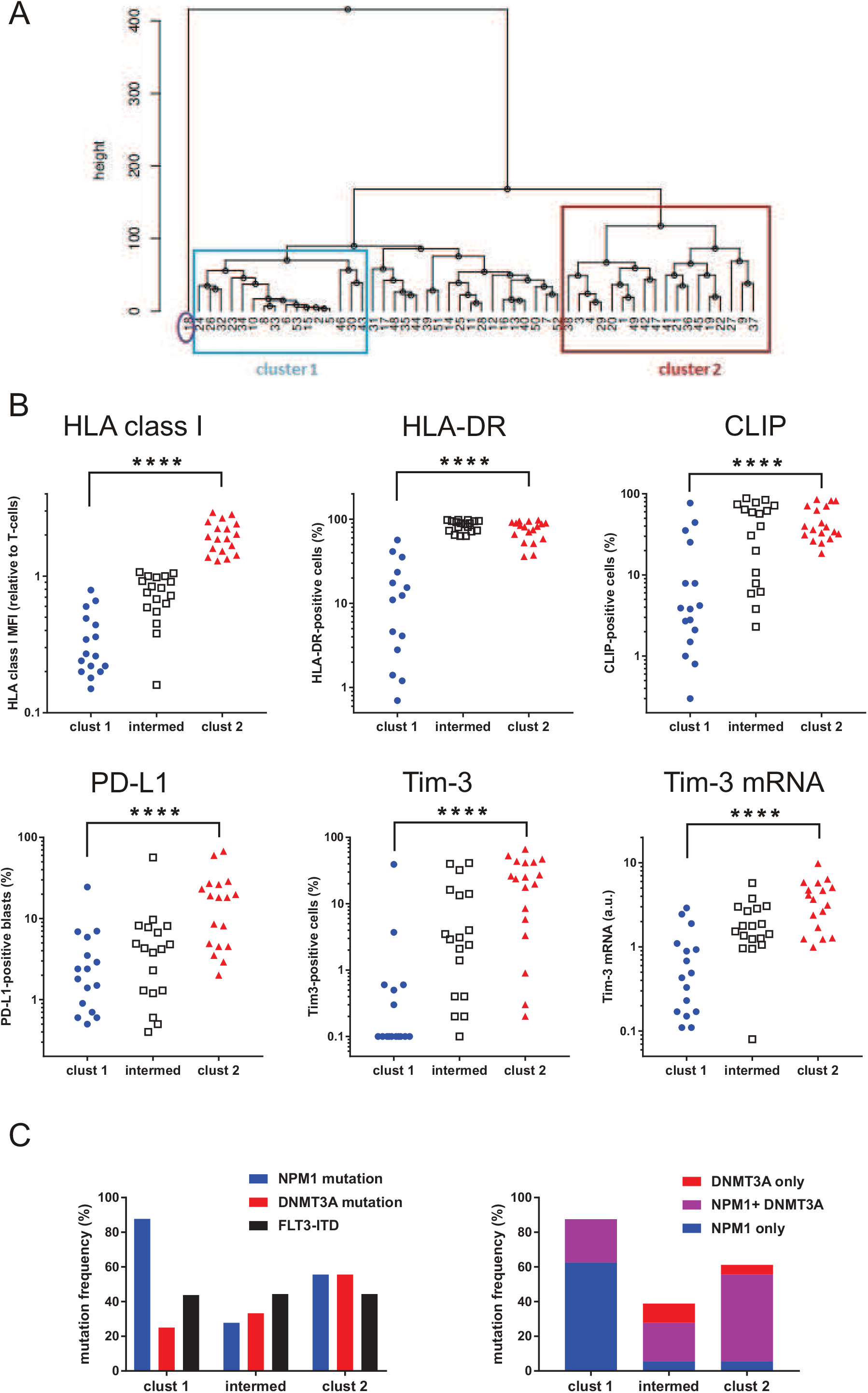
Results of hierarchical clustering of AML samples according to immune escape-related markers. A: Surface amounts of HLA class I, HLA-DR, CLIP, PD-L1, and TIM-3 were measured using flow cytometry and used as the input for clustering analysis. B: The graphs show the individual values in samples from the two clusters (cluster 1, cluster 2) defined in the tree diagram (top). The group denoted as "intermed" includes all the remaining samples except for the outlying ID 18. TIM-3 mRNA values were not used for hierarchical clustering, but are shown for comparison. Mann-Whitney test was performed to evaluate differences between cluster 1 and cluster 2, and p-value was found to be less than 0.0001 for all of the parameters. C: Relative incidence of recurrent mutations in the above defined groups (N = 16, 18, and 18 for cluster 1, intermediate group, and cluster 2, respectively).

Fig. 1C shows the relative incidence of recurrent mutations in the three groups resulting from the clustering analysis. The cluster 1 (N = 16) was characterized by a high percentage of NPM1 mutations and low incidence of DNMT3A mutations (Fig. 1C, left). In the cluster 2 (N = 18), a half of samples had both NPM1 and DNMT3A mutation (Fig. 1C, right). The occurrence of FLT3-ITD was closely similar in all the three groups.

### Impact of recurrent mutations on the immunophenotype

The whole cohort was then subdivided according to the presence of mutations in NPM1, FLT3, or DNMT3A, and the flow cytometry results were statistically evaluated. The measured values were usually not normally distributed, and we thus used the Mann-Whitney non-parametric test to assess differences between groups. As it is shown in Fig. 2 and Table 2, several statistically significant differences were found in this analysis:

**Table 2:** Summary of p-values obtained from statistical evaluation of the data shown in Fig. 2. The measured values of the indicated markers (left column) were compared in patient groups formed according to the presence of NPM1 mutation, FLT3-ITD, or DNMT3A mutation. CD47 values were compared using two-tailed t-test. The other parameters were not normally distributed and Mann-Whitney test was used for evaluation of differences between groups.

|  | NPM1 | FLT3-ITD | DNMT3A |
| --- | --- | --- | --- |
| HLA class I | 0.2126 | 0.5137 | 0.1228 |
| HLA-DR | * 0.0225 | 0.9039 | 0.5270 |
| CLIP | 0.8079 | * 0.0320 | * 0.0132 |
| PD-L1 | 0.8355 | 0.1243 | * 0.027 |
| Tim-3 | 0.1533 | 0.6196 | 0.2118 |
| CD47 | 0.0634 | 0.2440 | 0.1912 |

**Figure 2:**
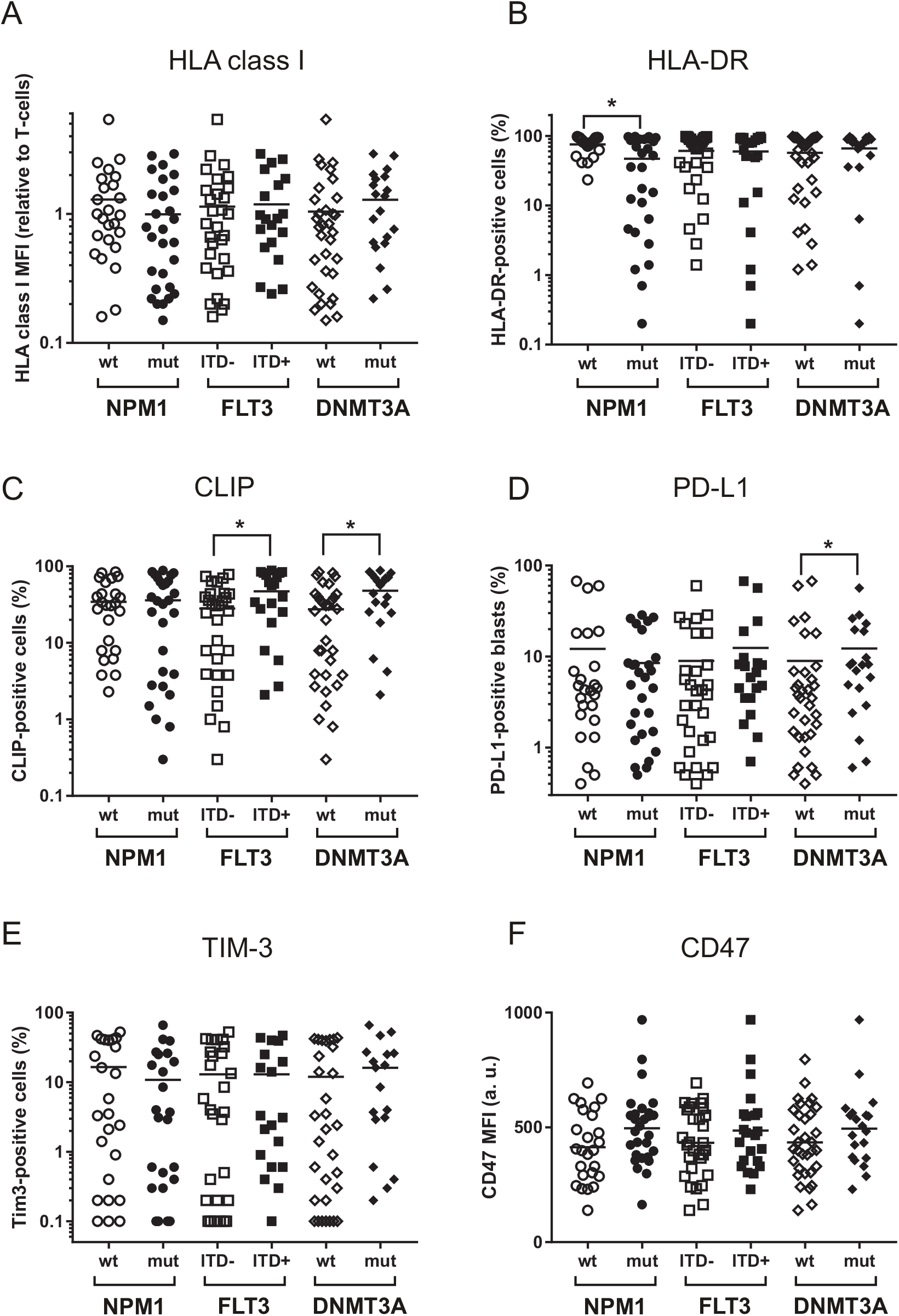
Comparison of the measured values in groups according to NPM1, FLT3, or DNMT3A mutation status. The cohort was subdivided according to NPM1 mutation/FLT3-ITD/DNMT3A mutation, and the values of the indicated surface marker were compared using the non-parametric Mann-Whitney test. All the resulting p-values are given in Table 2. The bars show the means, the star symbol denotes statistically significant difference between groups.

i. NPM1 mutation was associated with decreased HLA-DR expression (Fig. 2B, p = 0.0225).
ii. Patients with FLT3-ITD had higher surface amount of the invariant HLA-associated peptide CLIP (Fig. 2C, p = 0.032).
iii. Patients with DNMT3A mutations had higher levels of all the tested markers, the difference being statistically significant for CLIP (p = 0.0132) and PD-L1 (p = 0.027). On the other hand, the previously reported impact of FLT3-ITD on CD47 expression [16] was not confirmed in our cohort.

Characteristic features of the cluster 1 were decreased expression of HLA class I (the average MFI value for blasts was 33% of that for autologous T-cells) and low levels of other markers (Fig. 1B). This cluster included in particular many patients with NPM1 mutation and without concomitant DNMT3A mutation (Fig. 1C). On the other hand, the cluster 2 was enriched in samples with combined NPM1 and DNMT3 mutations. We thus stratified the whole cohort into four groups according to NPM1 and DNMT3A mutation status (Figure 3A). Isolated mutation in NPM1 was indeed associated with a decrease of HLA class I surface amount. Concomitant DNMT3A mutation prevented the effect of NPM1 mutation: the double-mutated group had similar HLA class I levels as the unmutated group. In the absence of DNMT3A mutation, the difference between NPM1 wild-type (wt) and NPM1 mutated (mut) samples was statistically significant (p = 0.0049 from Mann-Whitney test), as it was for the difference between DNMT3Awt and DNMT3Amut in the presence of NPM1 mutation (p = 0.0046).

**Figure 3:**
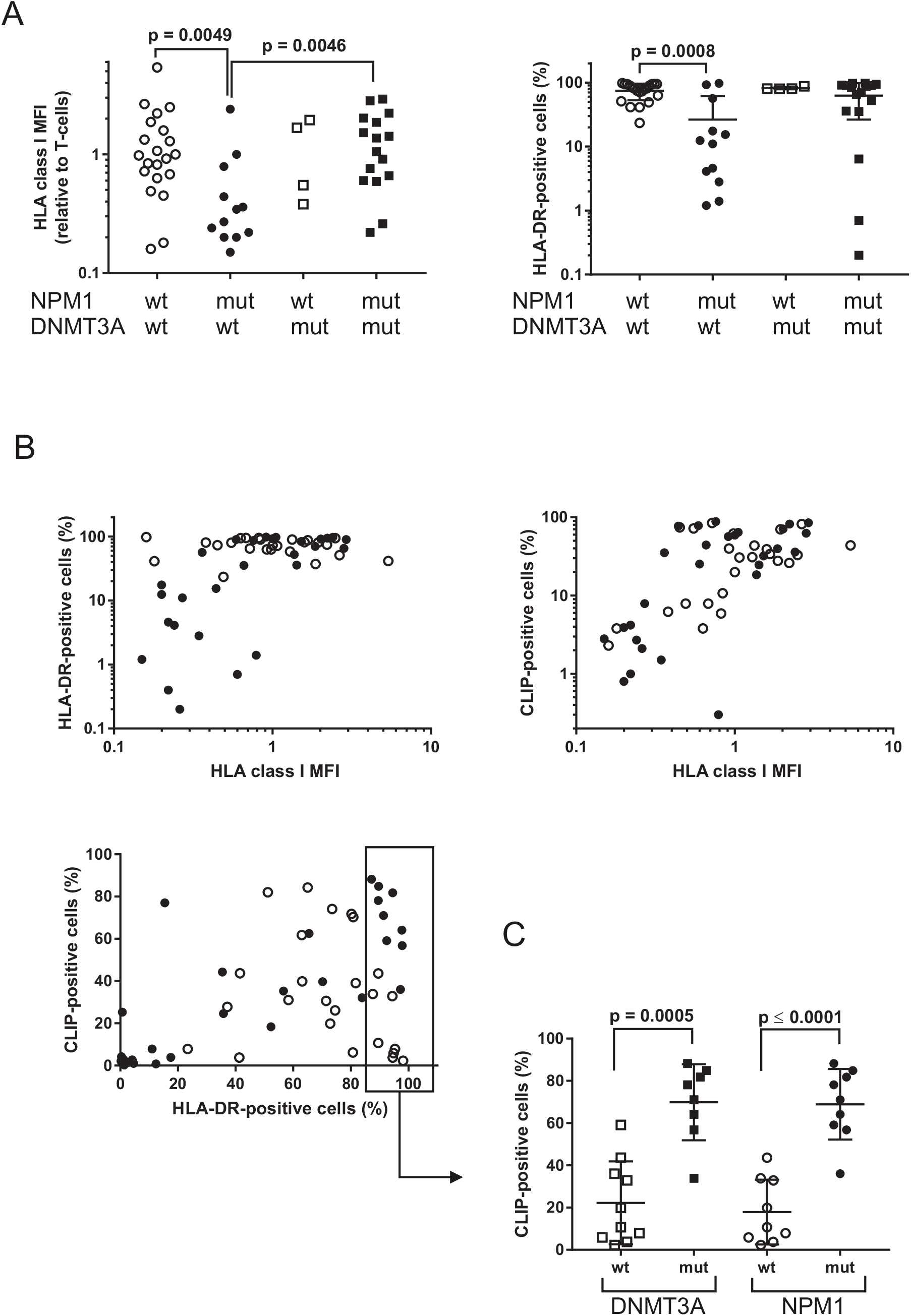
HLA-related markers in NPM1/DNMT3A groups. A: HLA class I and HLA-DR surface expression in patient subgroups according to NPM1 and DNMT3A mutations. Difference between groups were assessed using Mann-Whitney test, the resulting p-values are given in the graphs. B: Correlations between HLA class I, HLA-DR, and CLIP. Open symbols: samples with wild-type NPM1, closed symbols: samples with mutated NPM1. C: Samples with more than 85% HLA-DR-positive blasts were divided according to NPM1 or DNMT3A mutation and CLIP amounts were compared using the Mann-Whitney test. The resulting p-values are indicated in the graph.

HLA class I and HLA-DR are placed at the same DNA locus and their transcription is thus obviously correlated. Although the majority of AML blasts expressed HLA-DR, reduced amount of HLA-DR was observed in NPM1-mut group: p = 0.0225, resp. 0.0008 for comparison between NPM1-wt and NPM1-mut patients from the whole cohort (Fig. 2) or from DNMT3A-wt subgroup (Fig. 3A, right), respectively. Low HLA-DR expression usually correlated with low HLA class I expression (Fig. 3B). Nevertheless, two samples without NPM1 mutation had decreased HLA class I despite high HLA-DR levels (Fig. 3B, open circles indicate NPM1-wt cases, closed circles NPM1-mut).

As it was expected, samples with low HLA expression also had low amounts of the invariant peptide CLIP, which forms part of HLA class II (Fig. 3B). In samples with high HLA-DR levels, however, CLIP positivity was highly variable. We noted a marked difference between NPM1-wt and NPM1-mut samples with high HLA-DR expression: in the subset of samples with more than 85% HLA-DR positivity, the amount of CLIP was strongly associated with NPM1 and/or DNMT3A mutation (Fig. 3C, p = 0.0005 for comparison between DNMT3A-wt and DNMT3A-mut, p less than 0.0001 for comparison between NPM1-wt and NPM1-mut). In our cohort, DNMT3A mutation was mostly found in combination with NPM1 mutation (Fig. 1C, right) and the effects of these two mutations thus could not be separated.

### Transcript analysis

TIM-3-positive cell fraction determined by flow cytometry was not always well reproducible, maybe because this protein is removed from the cell surface by proteolytic cleavage [15]. Nevertheless, the surface TIM-3 positivity correlated with higher mRNA content (Fig. 4A). TIM-3 transcript also positively correlated with surface expression of PD-L1 (Fig. 4B) and CLIP (Fig. 4C). This indicates that the underlying mechanisms of immune escape are often activated simultaneously.

**Figure 4:**
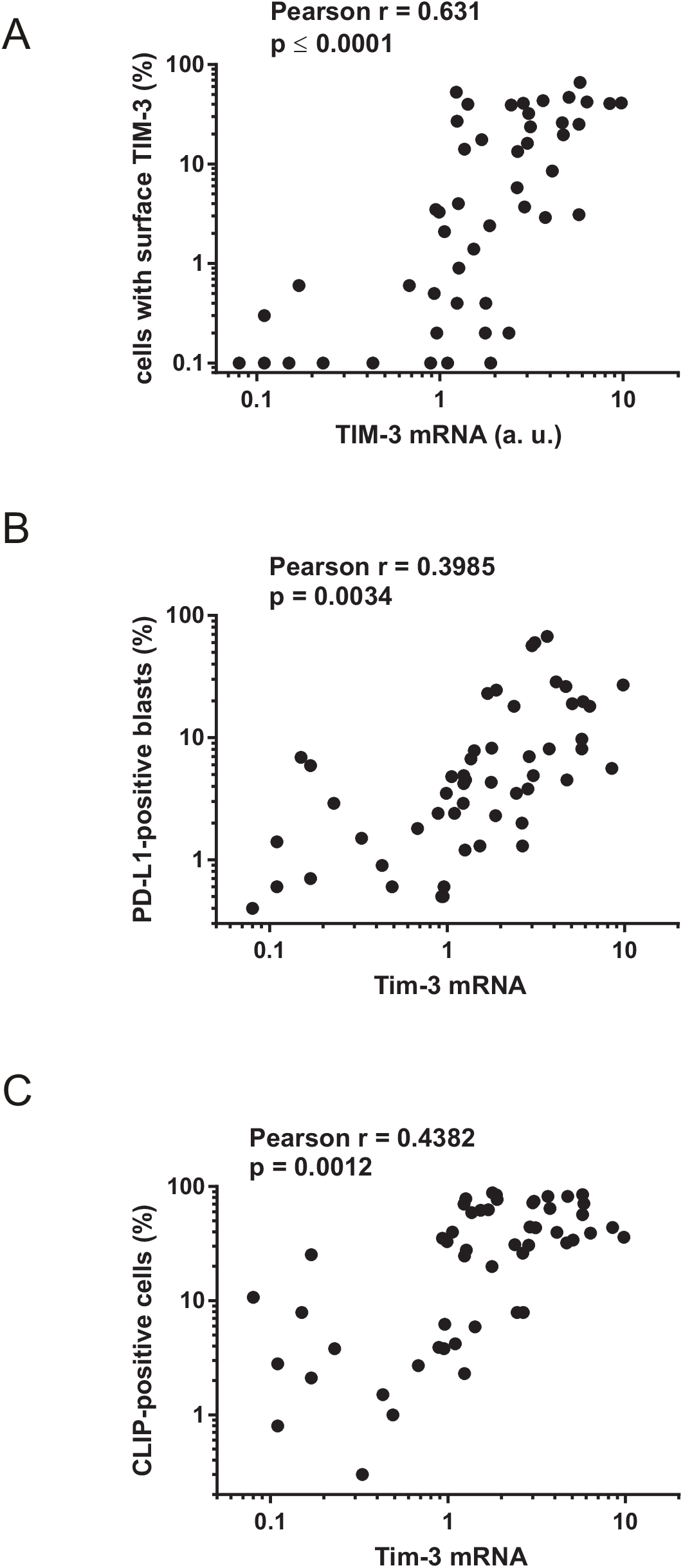
Correlation of TIM-3 mRNA with TIM-3 protein, PD-L1, and CLIP surface positivity of AML blasts. A: Correlation of TIM-3 trancript amount with the fraction of TIM-3-positive cells determined by flow cytometry in mononuclear cell preparations. B: The fraction of cells expressing the invariant CLIP peptide versus TIM-3 mRNA, C: The fraction of cells expressing the inhibitory receptor PD-L1 versus TIM-3 mRNA. Pearson correlation test was performed to assess linear correlation, the obtained correlation coefficients (r) and p-values are given in the graphs.

We have shown previously that the surface amount of PD-L1 correlates with the ratio of two transcription variants: the full-length v1 and a shorter v2 lacking the exon 2 [31,37]. Similar result was obtained in the present study (Fig. S2). Whereas the surface amount of PD-L1 did not display significant correlation with any of the individual transcript variants, the v1/v2 ratio correlated with the fraction of surface-positive cells. Interestingly, the positive correlation between surface protein and v1/v2 transcript was better in FLT3-ITD-positive samples (see the table in Fig. S2).

In the case of CD47, no correlation between transcript and protein was found (Figure S3).

As our patient cohort was relatively small, we analyzed an additional larger dataset obtained from the Vizome/BeatAML database, which contains transcriptomic data stratified according to different parameters [38]. The analysis confirmed transcript levels of all available HLA genes as well as that of TIM-3 (denoted as HAVCR2) to be significantly lower in the group with NPM1 mutation (N = 108) than in the group with wild-type NPM1 (N = 340). The graphs and p-values determined by standard two-tailed t-tests are given in Fig. S4.

### Survival analyses

Possible impact of blast immunophenotype on patient outcome was examined using standard survival analysis, and the resulting curves obtained for the overall survival (OS) and for relapse-free survival (RFS) are shown in Fig. 5. Patients with decreased HLA class I expression had significantly better outcome (p = 0.0407, resp. 0.0204 for OS, resp. RFS, Fig. 5A). This group mostly consists of patients with NPM1 mutation and without DNMT3A mutation, which also displayed low levels of the other markers of immune escape. In agreement with the fact that CLIP, PD-L1, and TIM-3 were to some extent correlated (Fig. 4), their impact on the survival was similar, high levels being always associated with worse prognosis (Fig. 5B-D). It seems likely that the regulation of different mechanisms enabling leukemia cells to escape from the immune system is to some extent shared, so that a resistant cell acquires a complex immunoresistant phenotype including increased expression of inhibitory receptors, reduced antigen presentation, and secretion of inhibitory molecules like TIM-3. Such phenotype would then be associated with unfavorable prognosis, and TIM-3 transcript could also be used as a prognostic factor reflecting this complex immune resistance (Fig. S5). We have shown previously that PD-L1 can also be measured as a ratio of v1/v2 transcript variants [37]. Indeed, similar results were obtained when v1/v2 mRNA PD-L1 levels were used for patient stratification, although the predictive power of the surface protein amount was better compared to the trancript (cf Fig. S6 and Fig. 5B).

**Figure 5:**
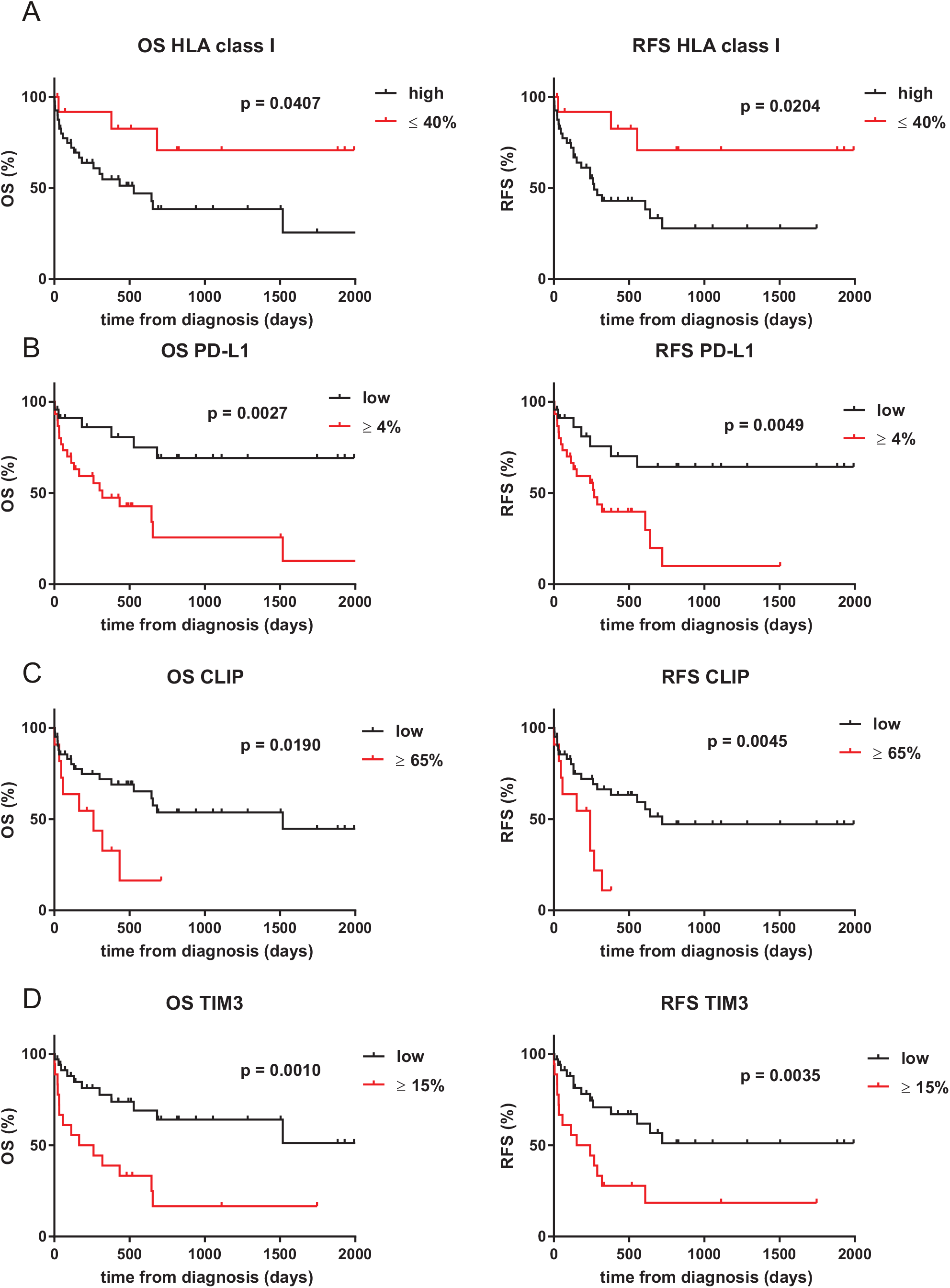
Survival analysis in groups according to selected immunophenotype markers. The analyses of the overall survival (OS, left column) and of the relapse-free survival (RFS, right column) in groups with low versus high levels of the indicated marker were performed using GraphPad Prism software, the obtained p-values for survival difference between groups are given in the graphs. The cutoff values were optimized with regard to the resulting p-values for OS.

### NK cell cytotoxic assay

Reduced HLA class I expression on leukemia cells could enhance the activity of natural killer (NK) cells, which are inhibited through binding of their inhibitory receptors (KIR) to HLA-C molecules on the target cell. We thus tested if the observed decrease of HLA expression could increase leukemia cell sensitivity to lysis by NK cells from a healthy donor. The effector cells used in the experiment had both 2DL1 and 2DL2 inhibitory KIR genes and were thus supposed to be inhibited by any HLA-C allele expressed on the target cells. NK cell-mediated killing of the positive control, i.e. K562 cells with very low HLA class I expression, occurred with 94% efficiency after 4h co-incubation at the effector:target ratio 10:1. On the other hand, the cytotoxicity observed in all samples of primary AML cells was limited (4 to 30%), and no difference was found between samples with low versus normal HLA class I, although we noted a trend to higher average efficiency of NK cells towards AML cells with low HLA amounts (Fig. 6).

**Figure 6:**
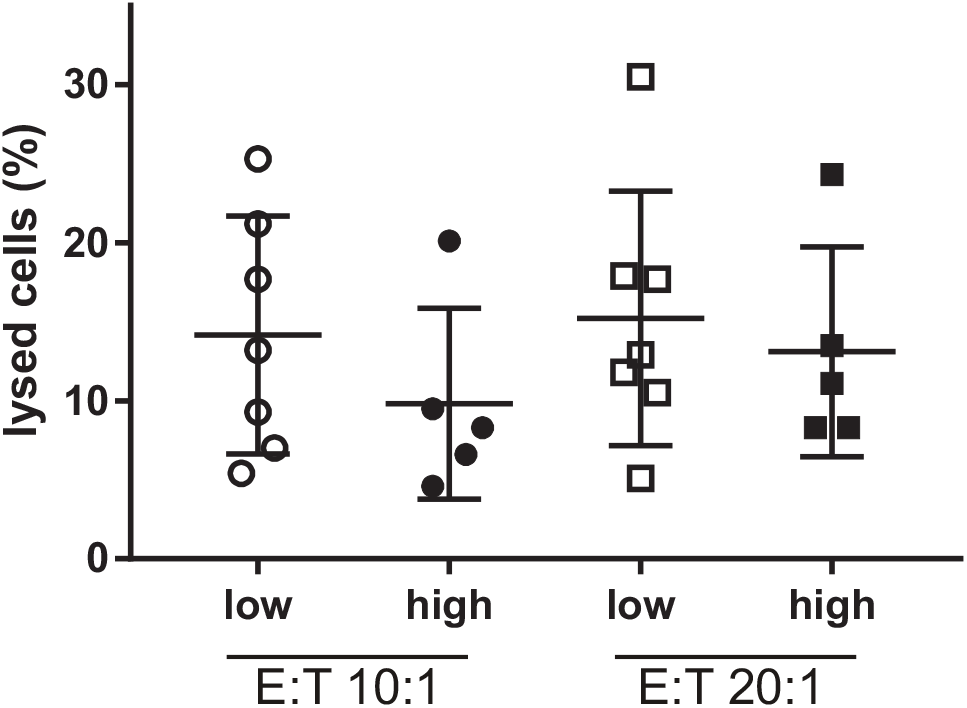
NK cell-mediated lysis of primary AML cells with low or high HLA class I expression. The cytotoxic assay with NK cells from a healthy donor was performed in triplicates at two different effector to target (E:T) ratios as indicated. The means of the triplicates are given in the Figure for samples with low HLA levels (N = 7, open symbols) or with normal HLA levels (N = 5, closed symbols). The bars indicate means and s.d. of the values shown in each group. In the positive control (K562 cells, not shown in the graph), the percentage of lysed cells reached 94% under the same experimental conditions at E:T ratio 10:1.

## Discussion

In this study, we analyzed selected markers of possible immune escape on AML blasts at diagnosis: HLA class I and HLA-DR as markers of decreased HLA expression, CLIP as a marker of reduced antigen presentation, the inhibitory ligand PD-L1 preventing T-cell cytotoxicity, TIM-3 as a marker of secretion of immunosuppressive molecules, and CD47 as an inhibitory ligand for macrophages. The majority of samples expressed at least one of these markers. The study was performed using samples obtained from leukapheresis, and was thus limited to patients with hyperleukocytosis, which are indicated to leukapheresis in order to reduce the tumor burden prior to chemotherapy. Therefore, it is possible that the cohort was enriched in cases with immune system suppression, which could allow for accumulation of high numbers of leukemia cells without excessive inflammation. Nevertheless, our results show that the mechanisms of immune escape known from solid tumors are also relevant for AML.

Fig. 2A,B and 3A suggest that HLA surface expression is reduced in association with NPM1 mutation. Decreased HLA-DR expression in AML with NPM1 mutation has already been reported previously [27], and HLA-DR positivity in this context was associated with worse survival [28]. Our results show that HLA class I follows similar expression pattern as HLA-DR (Fig. 3B), suggesting a lower transcription rate of the whole DNA locus in the presence of NPM1 mutation. This is supported by data available from Vizome/BeatAML database, which confirmed lower levels of HLA transcripts in samples with NPM1 mutation compared to samples with wild-type NPM1 (Fig. S4A). The effect of NPM1 mutation was reversed by DNMT3A mutation (Fig. 3A), which results in decreased DNMT3A activity. This is in agreement with a previous study describing enhanced HLA class I expression in neuronal cells with DNMT3A and DNMT1 knock-out [39].

Lower HLA class I expression could help leukemia cell escape from elimination by cytotoxic T-cells. We have previously reported that several HLA class I alleles were underrepresented in patients with NPM1 mutation [19,20]. This infers the possibility that NPM1-derived antigens are recognized by cytotoxic T-cells, which would create a selection pressure favoring the survival of cells with low HLA surface amount. Cells lacking HLA class I should be eliminated by NK cells due to the loss of inhibitory interaction between KIR and HLA-C. However, HLA class I expression on AML blasts was always partially preserved (at least 20% compared to the autologous T-lymphocytes, Fig. 2), and no significant increase in cell lysis by donor NK cells was observed in the group with low HLA levels (Fig. 6). The observed reduced HLA class I levels thus might represent a compromise between reduced sensitivity to T-cell attack and increased risk of lysis by NK cells, which would be reached during leukemia immunoediting. It appears that this phenotype is not associated with activation of other mechanisms of immune escape (Fig. 1B, cluster 1). According to survival analysis, the cases with decreased HLA expression have relatively good prognosis (Fig. 5A).

The invariant peptide CLIP is produced as a part of HLA class II and is replaced by antigens in the endoplasmic reticulum. In case of defective antigen presentation, CLIP is not removed from HLA molecules and is exposed on the cell surface. As we expected, samples with low HLA expression usually also had lower surface amounts of CLIP (Fig. 3B). However, in samples with high HLA-DR expression, high CLIP levels were significantly associated with NPM1 and/or DNMT3A mutations. This indicates lower efficiency in exposing specific antigens on HLA class II molecules in these AML subgroups. Reduced antigen presentation on HLA-DR is thus a relevant immune escape mechanism in AML, and seems to be specifically associated with NPM1 and/or DNMT3A mutation. Higher CLIP positivity was also found in samples with FLT3-ITD (Fig. 2C).

PD-L1 and PD-L2 are ligands to the inhibitory receptor PD-1 on T-cells, and their binding to PD-1 prevents the target cell lysis. Whereas PD-L2 was not detected on AML blasts, high surface PD-L1 positivity was associated with worse prognosis (Fig. 5B). In agreement with our previous study [31], PD-L1 surface protein levels correlated with v1/v2 PD-L1 mRNA (Fig. S2). In the previous study, the negative impact of high v1/v2 PD-L1 mRNA was restricted to patients with FLT3-ITD [31]. This could be due to the fact that the correlation between protein levels and v1/v2 transcript is clearly better in FLT3-ITD-positive cells (Fig. S2).

NPM1 was reported to be required for PD-L1 expression and NPM1 mutation thus could result in lower PD-L1 amounts. The average PD-L1 positivity in the group with mutated NPM1 was indeed slightly lower (Fig. 2), but the difference was not statistically significant (Table 2). On the other hand, analysis of the dataset from Vizome/BeatAML database showed a statistically significant decrease in TIM-3 transcript levels in association with NPM1 mutation (Fig. S4B).

The functionality of TIM-3 is highly context-dependent. TIM-3 positivity was associated with T-cell exhaustion, but also with increased cytotoxic capacity of NK cells [40,41]. In AML, higher numbers of TIM-3+ NK cells at diagnosis were recently reported to predict better outcome [42]. TIM-3 is also known to be present on the surface of AML blasts [43], but its role in leukemogenesis or in leukemia maintenance is still not clear. In its secreted form, it might assure distant inhibition of immune cells that are not in a direct contact with leukemia cells [15]. The observed correlation of PD-L1, and also of CLIP with TIM-3 mRNA (Fig. 4B,C) indicate that TIM-3 can indeed be a marker of immune resistance and that different mechanisms of immune escape may be activated in parallel.

DNMT3A mutation was associated with an increase in virtually all markers of immune escape (Fig. 2), and this fact may contribute to the known negative prognostic impact of this mutation. Correlation between lower expression/activity of DNMT3A and higher expression of PD-L1 was also observed in different tumor types [34–36].

CD47 is a protein associated with integrins and binds to an inhibitory receptor on macrophages. Its expression is transiently elevated during hematopoetic stem cell mobilization and release into the peripheral blood [44]. CD47 was detected in all samples, the mean amount being slightly higher in samples with NPM1 mutation (Fig. 2). This is in agreement with a previous study showing that CD47 positivity of AML blasts in the bone marrow correlated with NPM1 mutation [45]. However, the difference in MFI between NPM1-wt and NPM1-mut groups was not statistically significant in our cohort, and possible impact on the sensitivity of AML cells to an immune attack is not evident. Hierarchical clustering revealed one outlying sample (ID 18) with high TIM-3 and HLA class I expression (Fig. 1). The TIM-3 gene is located on the chromosome 5 and the aberrant mRNA TIM-3 level could thus be due to chromosome 5 duplication, which was found in the karyotype of this patient.

## Conclusions

At least one marker of possible immune escape was identified on leukemia blasts in a large part of AML patients with hyperleukocytosis. In the absence of DNMT3A mutation, samples with mutated NPM1 displayed decreased HLA expression, while they were usually negative in other markers. On the other hand, DNMT3A mutation was associated with higher levels of all the tested markers. High CLIP surface amount suggesting reduced antigen presentation was frequently found in patients with FLT3-ITD. CLIP, PD-L1, and TIM-3 were often co-expressed, suggesting parallel activation of different immune escape mechanisms. Our results indicate that the blast genotype is to some extent related to the immunophenotype, and the predictive value of particular mutations might also reflect an inherent cell resistance to the immune system.

## Methods

### Material

Primary cells from the peripheral blood of AML patients with hyperleukocytosis were obtained by leukapheresis at diagnosis, before therapy initiation. The leukapheretic products were diluted 20-fold in phosphate buffered saline (PBS) and the mononuclear cell fraction was then separated using Histopaque-1077 (Sigma, #H8889). The cells were then resuspended in RPMI 1640 medium with 10 % fetal calf serum and with antibiotics (100 U/ml penicillin, 100 μg/ml streptomycin), and aliquots were used for analysis of surface markers by flow cytometry and for mRNA isolation.

The analyses of karyotype and of mutations in NPM1, FLT3, and DNMT3A were performed as a part of routine clinical procedures.

The antibodies used were the following: CD45-V450 (#560367), CD4-BUV395 (#564724), CD8-BUV395 (#563795), and CD19-BUV737 (#564303) from BD Biosciences; HLA-DR-FITC (#11-9952-42), TIM-3-APC (#17-3109-42), CD47-APC (#17-0479-42), and PD-L2-APC (#17-5888-42) from eBioscience; CLIP-PE (sc-12725 PE) from Santa Cruz; PD-L1-PE (#1P-177-T100) from Exbio (Prague, Czech Republic). HLA class I antibody (ab2217) was purchased from Abcam and in house conjugated using the Lightning-Link Fluorescein Conjugation kit (#707-0010, Innova Biosciences).

### Flow cytometry

The mononuclear cells isolated from leukapheretic products were washed once in PBS, four tubes containing 1 milion cells in 50 μl PBS were prepared, antibodies were added (2 μl of each) and the samples were incubated for 30 min at 5°C. At the end of incubation, the cells were washed once in PBS (300g/5min/4°C), resuspended in ice-cold PBS, placed on ice, and immediately analyzed on a BD Fortessa flow cytometer using Application Settings. The settings were maintained using calibration beads. All the tubes contained CD45, CD4+CD8, and CD19 to discriminate blasts, T-cells, and B-cells. Leukemia blasts and lymphocytes were gated in CD45/SSC dotplots. The lymphocytes were further specified as T-cells using CD4 and CD8, or B-cells using CD19 positivity. No other antibody was added to the first tube, which was used to determine the background fluorescence. The background was subtracted from the mean fluorescence values (MFI) in each channel, for each subpopulation separately.

To quantify HLA class I expression, we used the mean fluorescence intensity (MFI) values. MFI for blasts were comparable with MFI for T-cells, whereas the values for B-cells were usually substantially higher (Supplementary Figure S1). As the surface amount of HLA class I and/or the affinity of the pan-HLA class I antibody might vary among individual HLA alleles, we expressed HLA class I values on blasts as relative to T-cells from the same sample. MFI was also used to quantify CD47 surface amount because the blast populations were rather homogenous with regard to CD47 expression. For the remaining markers, only a fraction of cells was usually positive, and these markers were quantified using the positive cell fraction (%) values.

### Real-time PCR

RNeasy Mini Kit (Qiagen) was used for RNA isolation from 2×10^7^ cells, and cDNA was generated by the reverse transcription on CFX96 real-time system (BioRad) using SensiFAST cDNA Synthesis Kit (Bioline). The quality and concentration of template RNA and of cDNA were assessed with NanoDrop One^C^ Microvolume UV-Vis Spectrophotometer (ThermoFisher Scientific). The relative amount of mRNA transcripts was measured by real-time PCR using SensiFAST SYBR N-ROX Kit (Bioline) and calculated by Bio-Rad CFX Manager Software. Primers for the individual genes (Table 3) were designed using PrimerBLAST software. For the relative quantification by 2^−ΔΔCt^ method, GAPDH expression was measured as a reference, using GAAACTGTGGCGTGATGGC and CCGTTCAGCTCAGGGATGAC as the forward and reverse primer, respectively.

**Table 3:**
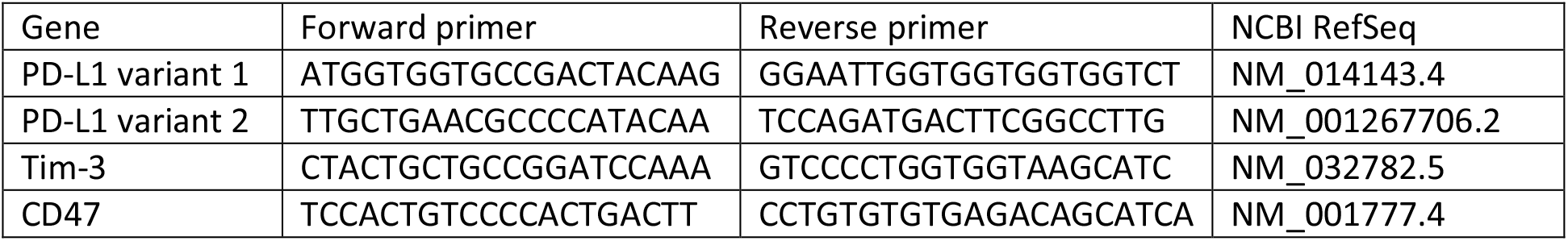
Primer sequences

### Test of natural killer (NK) cell cytotoxicity

The sensitivity of AML primary cells to lysis by NK cells was tested using cryopreserved samples of mononuclear cells obtained from leukapheresis as described above. A total of 12 samples included one group with low HLA class I expression (N = 7) and one group with high HLA class I expression (N = 5). HLA expression level was checked on defrozen samples and was found to be closely similar to that measured on fresh samples. K562 cells were used as a positive control. Although the target cell samples also contained other cells, the percentage of leukemia blasts was always high (e.g. in the group with low HLA expression, the mean percentage of blasts was 87%, range 76-96%, as determined from CD45/SSC dotplots).

NK cells were isolated and expanded from the buffy coat of a healthy donor. NK cells were separated from 1×10^8^ PBMC using NK Cell Isolation Kit (130-092-657, Miltenyi Biotech) and MACS column in the magnetic field (130-042-401, Miltenyi Biotech). They were expanded two weeks in CellGro GMP SCGM (20802 Cell Genix) with 5% inactivated human AB serum (AK9340, Akron Biotech), hIL-2 (1000 U/ml) (Proleukin S, Novartis), hIL-15 (20ng/ml)(130-095-765, Miltenyi Biotech) and gentamycin (40ug/ml)(G1272, Sigma). Cells were stimulated with Activation/Expansion Kit (130-094-483, Miltenyi Biotech) consisting of Bead Particles loaded with NKp46 and CD2 antibodies. The KIR genotype of NK cells included 2DL1, 2DL2, 2DL3, 2DL4, 2DL5, 2DS1, 2DS2, 2DS3, 2DS5, 3DL2, 3DL3, 3DS1, 2DP1 and 3DP1 genes.

Target cells were stained with carboxyfluorescein diacetate succinimidyl ester (CFSE, 565082 BD Biosciences) and mixed with the effector NK cells in 1:10 or 1:20 ratio. The experiment was performed in triplets with 10 000 target cells per well (96-well plate). Wells without effector NK cells were used to determine the fraction of spontaneously dead cells (% spont dead). The plate was incubated for 4h at 37°C (in a CO_2_ incubator). Then, the CFSE+ cell viability was assessed by flow-cytometry using propidium iodide staining. The cell debris were outgated from scattergrams, the singlet events were selected from FSC-A vs FSC-H dotplots and CFSE+ target cells were then gated from CFSE vs SSC-A dotplots. The cytotoxicity was calculated from the following formula: % cytotoxicity = (% dead in the sample with NK - % spont dead)/(100 - % spont dead)

### Data analysis

Hierarchical clustering was performed using the McQuitty clustering method in the free Wessa software (https://www.wessa.net). GraphPad Prism software (v.7) was used for statistical evaluation of the data (t-test, Mann-Whitney test, Pearson correlation) and for survival analyses. The p-value limit for significant differences between groups was set to 0.05.

## Funding Information

This work was financially supported by the European Regional Development Fund, by the state budget of the Czech Republic (project AIIHHP: CZ.02.1.01/0.0/0.0/16_025/0007428, OP RDE, Ministry of Education, Youth and Sports), and by the Ministry of Health of the Czech Republic (project for conceptual development of the research organization No 00023736).

## Acknowledgement

The authors acknowledge the expert technical assistance provided by P.Otevřelová. Information about NPM1 mutation type was provided by E.Cerovská.

## Author contributions

Conceptualization, K:K. and B.B.; Methodology, K.K. and B.B.; Investigation, K.K., B.B., J.M., M.P., Š.R.; Resources, Z.G.; Data Curation, K.K., B.B., J.M., Š.R., and C.Š.; Writing – Original Draft Preparation, K.K.; Writing – Review & Editing, B.B., J.M., M.P., Š.R., C.Š., Z.G.; Funding Acquisition, K.K.

## Conflict of Interests

The authors declare no conflict of interest. The sponsors had no role in the design, execution, interpretation, or writing of the study.

## Ethics Statement

All patients provided their written informed consent as to the use of their biological material for research purposes. The project was approved by the Ethics Committee of the Institute of Hematology in June 2015. All procedures followed were in accordance with the ethical standards of the responsible committees on human experimentation (institutional and national) and with the Helsinki Declaration of 1975, as revised in 2008.

## Notes

### Competing Interest Statement

The authors have declared no competing interest.

## References

[1] Buggins, A.G.; Milojkovic, D.; Arno, M.J.; Lea, N.C.; Mufti, G.J.; Thomas, N.S.; Hirst, W.J. Microenvironment Produced by Acute Myeloid Leukemia Cells Prevents T Cell Activation and Proliferation by Inhibition of NF-kappaB, C-Myc, and pRb Pathways. J. Immunol. 2001, 167, 6021–6030.

[2] Knaus, H.A.; Berglund, S.; Hackl, H.; Blackford, A.L.; Zeidner, J.F.; Montiel-Esparza, R.; Mukhopadhyay, R.; Vanura, K.; Blazar, B.R.; Karp, J.E. et al. Signatures of CD8+ T Cell Dysfunction in AML Patients and their Reversibility with Response to Chemotherapy. JCI Insight 2018, 3, e120974. doi: 10.1172/jci.insight.120974.

[3] Le Dieu, R.; Taussig, D.C.; Ramsay, A.G.; Mitter, R.; Miraki-Moud, F.; Fatah, R.; Lee, A.M.; Lister, T.A.; Gribben, J.G. Peripheral Blood T Cells in Acute Myeloid Leukemia (AML) Patients at Diagnosis have Abnormal Phenotype and Genotype and Form Defective Immune Synapses with AML Blasts. Blood 2009, 114, 3909–3916.

[4] Lamble, A.J.; Kosaka, Y.; Laderas, T.; Maffit, A.; Kaempf, A.; Brady, L.K.; Wang, W.; Long, N.; Saultz, J.N.; Mori, M. et al. Reversible Suppression of T Cell Function in the Bone Marrow Microenvironment of Acute Myeloid Leukemia. Proc. Natl. Acad. Sci. U. S. A. 2020, 117, 14331–14341.

[5] Tan, J.; Yu, Z.; Huang, J.; Chen, Y.; Huang, S.; Yao, D.; Xu, L.; Lu, Y.; Chen, S.; Li, Y. Increased PD-1+Tim-3+ Exhausted T Cells in Bone Marrow may Influence the Clinical Outcome of Patients with AML. Biomark Res. 2020, 8, 6–8. eCollection 2020.

[6] Barrett, A.J. Acute Myeloid Leukaemia and the Immune System: Implications for Immunotherapy. Br. J. Haematol. 2020, 188, 147–158.

[7] Vago, L.; Gojo, I. Immune Escape and Immunotherapy of Acute Myeloid Leukemia. J. Clin. Invest. 2020, 130, 1552–1564.

[8] Valent, P.; Sadovnik, I.; Eisenwort, G.; Bauer, K.; Herrmann, H.; Gleixner, K.V.; Schulenburg, A.; Rabitsch, W.; Sperr, W.R.; Wolf, D. Immunotherapy-Based Targeting and Elimination of Leukemic Stem Cells in AML and CML. Int. J. Mol. Sci. 2019, 20, 4233. doi: 10.3390/ijms20174233.

[9] van Luijn, M.M.; van de Loosdrecht, A. A.; Lampen, M.H.; van Veelen, P.A.; Zevenbergen, A.; Kester, M.G.; de Ru, A.H.; Ossenkoppele, G.J.; van Hall, T.; van Ham, S.M. Promiscuous Binding of Invariant Chain-Derived CLIP Peptide to Distinct HLA-I Molecules Revealed in Leukemic Cells. PLoS One 2012, 7, e34649.

[10] van Luijn, M.M.; Chamuleau, M.E.; Ossenkoppele, G.J.; van de Loosdrecht, A. A.; Marieke van Ham, S. Tumor Immune Escape in Acute Myeloid Leukemia: Class II-Associated Invariant Chain Peptide Expression as Result of Deficient Antigen Presentation. Oncoimmunology 2012, 1, 211–213.

[11] van den Ancker, W.; van Luijn, M.M.; Chamuleau, M.E.; Kelder, A.; Feller, N.; Terwijn, M.; Zevenbergen, A.; Schuurhuis, G.J.; Ham, S.M.; Westers, T.M. et al. High Class II-Associated Invariant Chain Peptide Expression on Residual Leukemic Cells is Associated with Increased Relapse Risk in Acute Myeloid Leukemia. Leuk. Res. 2014, 38, 691–693.

[12] Monney, L.; Sabatos, C.A.; Gaglia, J.L.; Ryu, A.; Waldner, H.; Chernova, T.; Manning, S.; Greenfield, E.A.; Coyle, A.J.; Sobel, R.A. et al. Th1-Specific Cell Surface Protein Tim-3 Regulates Macrophage Activation and Severity of an Autoimmune Disease. Nature 2002, 415, 536–541.

[13] Jan, M.; Chao, M.P.; Cha, A.C.; Alizadeh, A.A.; Gentles, A.J.; Weissman, I.L.; Majeti, R. Prospective Separation of Normal and Leukemic Stem Cells Based on Differential Expression of TIM3, a Human Acute Myeloid Leukemia Stem Cell Marker. Proc. Natl. Acad. Sci. U. S. A. 2011, 108, 5009–5014.

[14] Kikushige, Y.; Miyamoto, T.; Yuda, J.; Jabbarzadeh-Tabrizi, S.; Shima, T.; Takayanagi, S.; Niiro, H.; Yurino, A.; Miyawaki, K.; Takenaka, K. et al. A TIM-3/Gal-9 Autocrine Stimulatory Loop Drives Self-Renewal of Human Myeloid Leukemia Stem Cells and Leukemic Progression. Cell. Stem Cell. 2015, 17, 341–352.

[15] Goncalves Silva, I.; Ruegg, L.; Gibbs, B.F.; Bardelli, M.; Fruehwirth, A.; Varani, L.; Berger, S.M.; Fasler-Kan, E.; Sumbayev, V.V. The Immune Receptor Tim-3 Acts as a Trafficker in a Tim-3/Galectin-9 Autocrine Loop in Human Myeloid Leukemia Cells. Oncoimmunology 2016, 5, e1195535.

[16] Majeti, R.; Chao, M.P.; Alizadeh, A.A.; Pang, W.W.; Jaiswal, S.; Gibbs, K.D.,Jr; van Rooijen, N.; Weissman, I.L. CD47 is an Adverse Prognostic Factor and Therapeutic Antibody Target on Human Acute Myeloid Leukemia Stem Cells. Cell 2009, 138, 286–299.

[17] Pietsch, E.C.; Dong, J.; Cardoso, R.; Zhang, X.; Chin, D.; Hawkins, R.; Dinh, T.; Zhou, M.; Strake, B.; Feng, P.H. et al. Anti-Leukemic Activity and Tolerability of Anti-Human CD47 Monoclonal Antibodies. Blood Cancer. J. 2017, 7, e536.

[18] Petrova, P.S.; Viller, N.N.; Wong, M.; Pang, X.; Lin, G.H.; Dodge, K.; Chai, V.; Chen, H.; Lee, V.; House, V. et al. TTI-621 (SIRPαFc): A CD47-Blocking Innate Immune Checkpoint Inhibitor with Broad Antitumor Activity and Minimal Erythrocyte Binding. Clin. Cancer Res. 2017, 23, 1068–1079.

[19] Kuzelova, K.; Brodska, B.; Fuchs, O.; Dobrovolna, M.; Soukup, P.; Cetkovsky, P. Altered HLA Class I Profile Associated with Type A/D Nucleophosmin Mutation Points to Possible Anti-Nucleophosmin Immune Response in Acute Myeloid Leukemia. PLoS One 2015, 10, e0127637.

[20] Kuzelova, K.; Brodska, B.; Schetelig, J.; Rollig, C.; Racil, Z.; Walz, J.S.; Helbig, G.; Fuchs, O.; Vrana, M.; Pecherkova, P. et al. Association of HLA Class I Type with Prevalence and Outcome of Patients with Acute Myeloid Leukemia and Mutated Nucleophosmin. PLoS One 2018, 13, e0204290.

[21] Narayan, R.; Olsson, N.; Wagar, L.E.; Medeiros, B.C.; Meyer, E.; Czerwinski, D.; Khodadoust, M.S.; Zhang, L.; Schultz, L.; Davis, M.M. et al. Acute Myeloid Leukemia Immunopeptidome Reveals HLA Presentation of Mutated Nucleophosmin. PLoS One 2019, 14, e0219547.

[22] Berlin, C.; Kowalewski, D.J.; Schuster, H.; Mirza, N.; Walz, S.; Handel, M.; Schmid-Horch, B.; Salih, H.R.; Kanz, L.; Rammensee, H.G. et al. Mapping the HLA Ligandome Landscape of Acute Myeloid Leukemia: A Targeted Approach Toward Peptide-Based Immunotherapy. Leukemia 2014.

[23] Greiner, J.; Schneider, V.; Schmitt, M.; Gotz, M.; Dohner, K.; Wiesneth, M.; Dohner, H.; Hofmann, S. Immune Responses Against the Mutated Region of Cytoplasmatic NPM1 might Contribute to the Favorable Clinical Outcome of AML Patients with NPM1 Mutations (NPM1mut). Blood 2013, 122, 1087–1088.

[24] Greiner, J.; Ono, Y.; Hofmann, S.; Schmitt, A.; Mehring, E.; Gotz, M.; Guillaume, P.; Dohner, K.; Mytilineos, J.; Dohner, H. et al. Mutated Regions of Nucleophosmin 1 Elicit both CD4(+) and CD8(+) T-Cell Responses in Patients with Acute Myeloid Leukemia. Blood 2012, 120, 1282–1289.

[25] van der Lee, D. I.; Reijmers, R.M.; Honders, M.W.; Hagedoorn, R.S.; de Jong, R.C.; Kester, M.G.; van der Steen, D. M.; de Ru, A.H.; Kweekel, C.; Bijen, H.M. et al. Mutated Nucleophosmin 1 as Immunotherapy Target in Acute Myeloid Leukemia. J. Clin. Invest. 2019, 129, 774–785.

[26] Forghieri, F.; Riva, G.; Lagreca, I.; Barozzi, P.; Vallerini, D.; Morselli, M.; Paolini, A.; Bresciani, P.; Colaci, E.; Maccaferri, M. et al. Characterization and Dynamics of Specific T Cells Against Nucleophosmin-1 (NPM1)-Mutated Peptides in Patients with NPM1-Mutated Acute Myeloid Leukemia. Oncotarget 2019, 10, 869–882.

[27] Chauhan, P.S.; Ihsan, R.; Singh, L.C.; Gupta, D.K.; Mittal, V.; Kapur, S. Mutation of NPM1 and FLT3 Genes in Acute Myeloid Leukemia and their Association with Clinical and Immunophenotypic Features. Dis. Markers 2013, 35, 581–588.

[28] Chen, C.Y.; Chou, W.C.; Tsay, W.; Tang, J.L.; Yao, M.; Huang, S.Y.; Tien, H.F. Hierarchical Cluster Analysis of Immunophenotype Classify AML Patients with NPM1 Gene Mutation into Two Groups with Distinct Prognosis. BMC Cancer 2013, 13, 107–107.

[29] Mason, E.F.; Kuo, F.C.; Hasserjian, R.P.; Seegmiller, A.C.; Pozdnyakova, O. A Distinct Immunophenotype Identifies a Subset of NPM1-Mutated AML with TET2 Or IDH1/2 Mutations and Improved Outcome. Am. J. Hematol. 2018, 93, 504–510.

[30] Qin, G.; Wang, X.; Ye, S.; Li, Y.; Chen, M.; Wang, S.; Qin, T.; Zhang, C.; Li, Y.; Long, Q. et al. NPM1 Upregulates the Transcription of PD-L1 and Suppresses T Cell Activity in Triple-Negative Breast Cancer. Nat. Commun. 2020, 11, 1669–z.

[31] Brodska, B.; Otevrelova, P.; Salek, C.; Fuchs, O.; Gasova, Z.; Kuzelova, K. High PD-L1 Expression Predicts for Worse Outcome of Leukemia Patients with Concomitant NPM1 and FLT3 Mutations. Int. J. Mol. Sci. 2019, 20, 10.3390/ijms20112823.

[32] Li, C.; Chen, X.; Yu, X.; Zhu, Y.; Ma, C.; Xia, R.; Ma, J.; Gu, C.; Ye, L.; Wu, D. Tim-3 is Highly Expressed in T Cells in Acute Myeloid Leukemia and Associated with Clinicopathological Prognostic Stratification. Int. J. Clin. Exp. Pathol. 2014, 7, 6880–6888.

[33] Zhang, Q.; Wu, X.; Cao, J.; Gao, F.; Huang, K. Association between Increased Mutation Rates in DNMT3A and FLT3-ITD and Poor Prognosis of Patients with Acute Myeloid Leukemia. Exp. Ther. Med. 2019, 18, 3117–3124.

[34] Li, X.; Wang, Z.; Huang, J.; Luo, H.; Zhu, S.; Yi, H.; Zheng, L.; Hu, B.; Yu, L.; Li, L. et al. Specific Zinc Finger-Induced Methylation of PD-L1 Promoter Inhibits its Expression. FEBS Open Bio 2019, 9, 1063–1070.

[35] Darvin, P.; Sasidharan Nair, V.; Elkord, E. PD-L1 Expression in Human Breast Cancer Stem Cells is Epigenetically Regulated through Posttranslational Histone Modifications. J. Oncol. 2019, 2019, 3958908.

[36] Chatterjee, A.; Rodger, E.J.; Ahn, A.; Stockwell, P.A.; Parry, M.; Motwani, J.; Gallagher, S.J.; Shklovskaya, E.; Tiffen, J.; Eccles, M.R. et al. Marked Global DNA Hypomethylation is Associated with Constitutive PD-L1 Expression in Melanoma. iScience 2018, 4, 312–325.

[37] Brodska, B.; Otevrelova, P.; Kuzelova, K. Correlation of PD-L1 Surface Expression on Leukemia Cells with the Ratio of PD-L1 mRNA Variants and with Electrophoretic Mobility. Cancer. Immunol. Res. 2016, 4, 815–819.

[38] Tyner, J.W.; Tognon, C.E.; Bottomly, D.; Wilmot, B.; Kurtz, S.E.; Savage, S.L.; Long, N.; Schultz, A.R.; Traer, E.; Abel, M. et al. Functional Genomic Landscape of Acute Myeloid Leukaemia. Nature 2018, 562, 526–531.

[39] Feng, J.; Zhou, Y.; Campbell, S.L.; Le, T.; Li, E.; Sweatt, J.D.; Silva, A.J.; Fan, G. Dnmt1 and Dnmt3a Maintain DNA Methylation and Regulate Synaptic Function in Adult Forebrain Neurons. Nat. Neurosci. 2010, 13, 423–430.

[40] Ndhlovu, L.C.; Lopez-Vergès, S.; Barbour, J.D.; Jones, R.B.; Jha, A.R.; Long, B.R.; Schoeffler, E.C.; Fujita, T.; Nixon, D.F.; Lanier, L.L. Tim-3 Marks Human Natural Killer Cell Maturation and Suppresses Cell-Mediated Cytotoxicity. Blood 2012, 119, 3734–3743.

[41] Gleason, M.K.; Lenvik, T.R.; McCullar, V.; Felices, M.; O’Brien, M.S.; Cooley, S.A.; Verneris, M.R.; Cichocki, F.; Holman, C.J.; Panoskaltsis-Mortari, A. et al. Tim-3 is an Inducible Human Natural Killer Cell Receptor that Enhances Interferon Gamma Production in Response to Galectin-9. Blood 2012, 119, 3064–3072.

[42] Rakova, J.; Truxova, I.; Holicek, P.; Salek, C.; Hensler, M.; Kasikova, L.; Pasulka, J.; Holubova, M.; Kovar, M.; Lysak, D. et al. TIM-3 Levels Correlate with Enhanced NK Cell Cytotoxicity and Improved Clinical Outcome in AML Patients. Oncoimmunology 2021, 10, 1889822.

[43] Kikushige, Y.; Miyamoto, T. TIM-3 as a Novel Therapeutic Target for Eradicating Acute Myelogenous Leukemia Stem Cells. Int. J. Hematol. 2013, 98, 627–633.

[44] Jaiswal, S.; Jamieson, C.H.; Pang, W.W.; Park, C.Y.; Chao, M.P.; Majeti, R.; Traver, D.; van Rooijen, N.; Weissman, I.L. CD47 is Upregulated on Circulating Hematopoietic Stem Cells and Leukemia Cells to Avoid Phagocytosis. Cell 2009, 138, 271–285.

[45] Galli, S.; Zlobec, I.; Schürch, C.; Perren, A.; Ochsenbein, A.F.; Banz, Y. CD47 Protein Expression in Acute Myeloid Leukemia: A Tissue Microarray-Based Analysis. Leuk. Res. 2015, 39, 749–756.

