## Supplementary Figures for "Associations between recurrent mutations and blast immunophenotype in acute myeloid leukemia"

Supplementary Material
Figures S1 to S6

Figure S1: HLA class I expression on AML blasts, T-cells, and B-cells

The mean fluorescence intensity (MFI) of HLA class I-FITC signal on blasts, T-cells, and B-cells. Blasts and lymphocytes were gated in CD45/SSC dotplots. Cells in the lymphocyte gate were further discriminated using CD4+CD8 or CD19-positivity to gate T-cells or B-cells, respectively. The points belonging to the same patient sample are connected with solid lines.

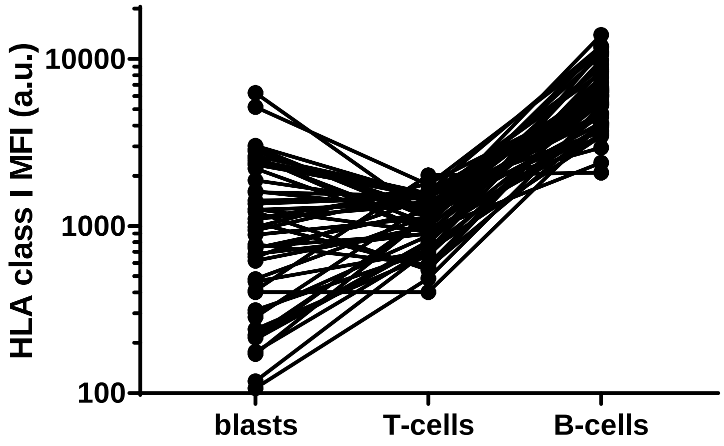

Figure S2: Assesment of PD-L1 transcript levels

Correlation of PD-L1 transcript variants v1 (A), v2 (B) or of their ratio (C) with the fraction of surface-positive cells. The transcript was assessed by RT-PCR from mRNA samples. Open circles: samples without FLT3-ITD, closed circles: samples with FLT3-ITD. Linear correlation was assessed using GraphPad Prism v.7, the Pearson correlation coefficient (r) and p-value are given in the table (bottom).

A. v1 = full-length variant

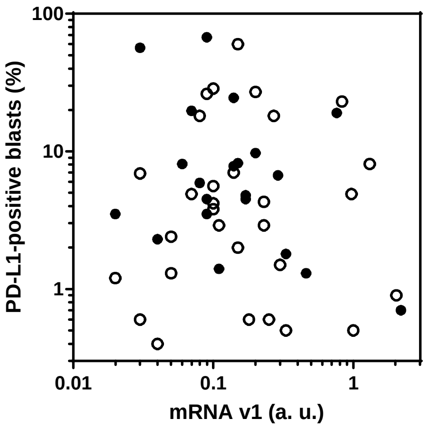


B. v2 = missing exon 2

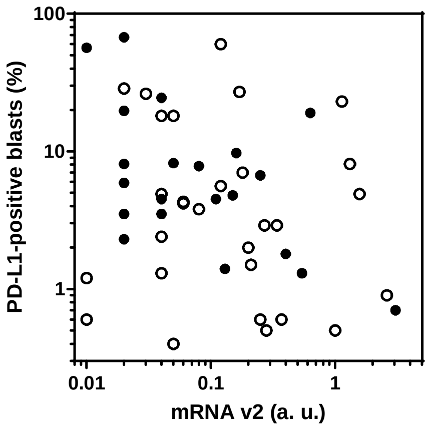


C. v1/v2 ratio

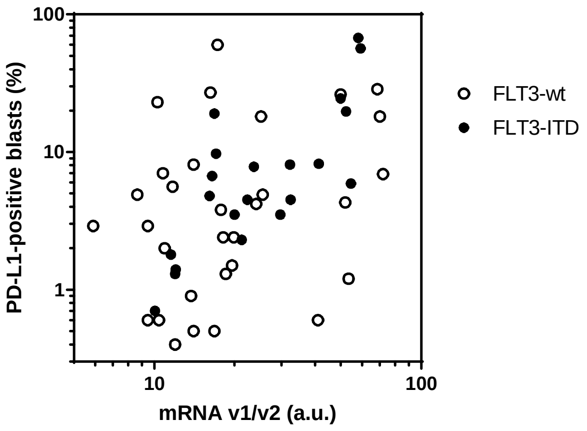


| | whole cohort | FLT3-WT | FLT3-ITD |
|---|---|---|---|
| v1 | r = -0.1375<br>p = 0.3312 | r = -0.085<br>p = 0.6506 | r = -0.187<br>p = 0.4169 |
| v2 | r = -0.1627<br>p = 0.2490 | r = -0.113<br>p = 0. 5449 | r = -0.2015<br>p = 0.3812 |
| v1/v2 | r = 0.4296<br>p = 0. 0015 | r = 0.2162<br>p = 0.2428 | r = 0.7106<br>p = 0.0003 |

Figure S3: Analysis of CD47 transcript

Correlation of CD47 transcript with the amount of surface protein measured as the mean fluorescence intensity of CD47-FITC signal.

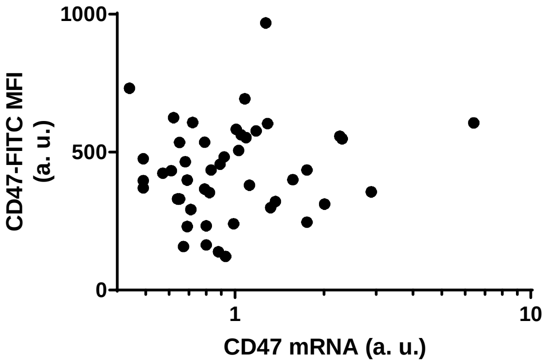

Figure S4: Amount of HLA and TIM-3 transcripts in primary AML cells – effect of NPM1 mutation

Transcriptomic data are available from www.vizome.org, in the BeatAML section. We compared the transcript levels of HLA (panel A) and of TIM-3 (denoted as HAVCR2 in the database) in samples with unmutated (NPMwt, N=340) and mutated (NPMmut, N=108) nucleophosmin 1 by two-tailed unpaired t-test, the resulting p-values are given in the graphs.

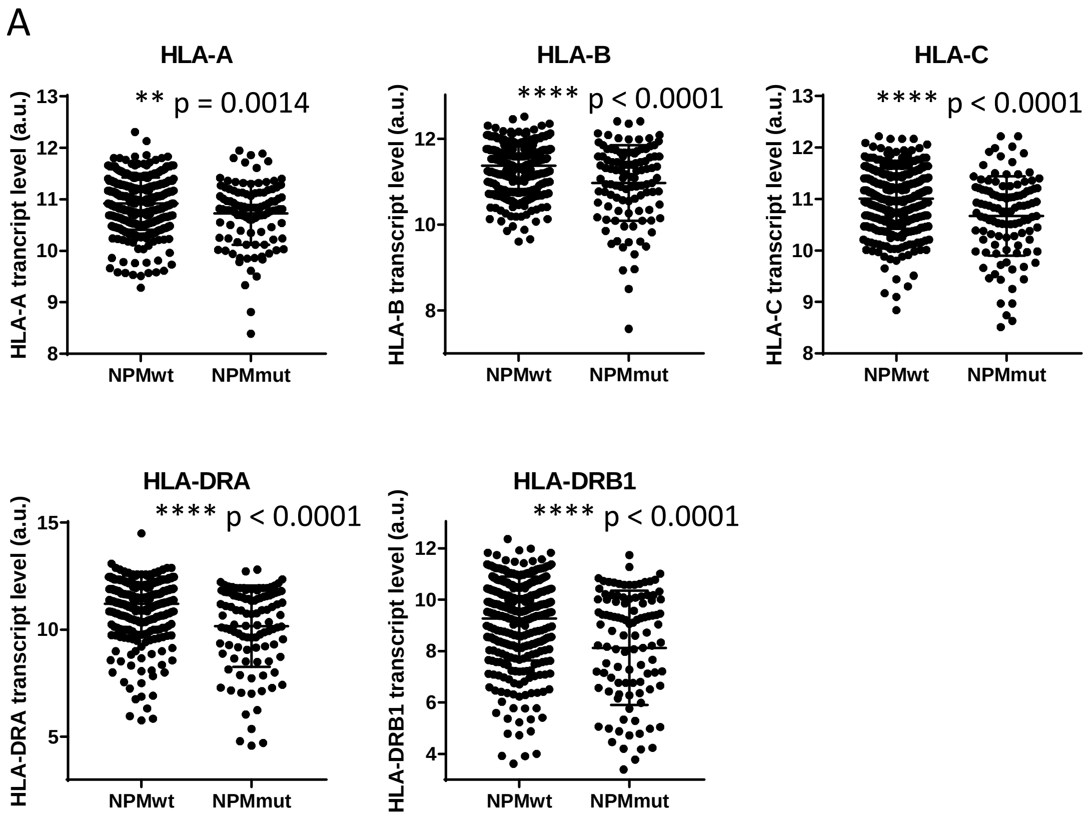


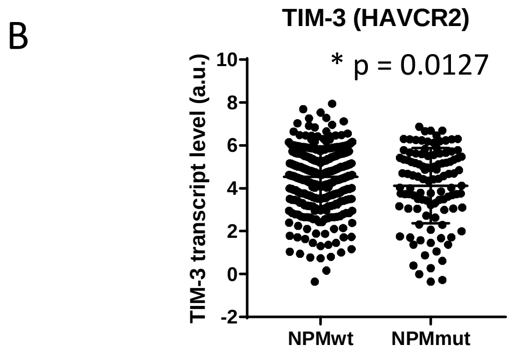

Figure S5: Impact of TIM-3 mRNA on patient survival

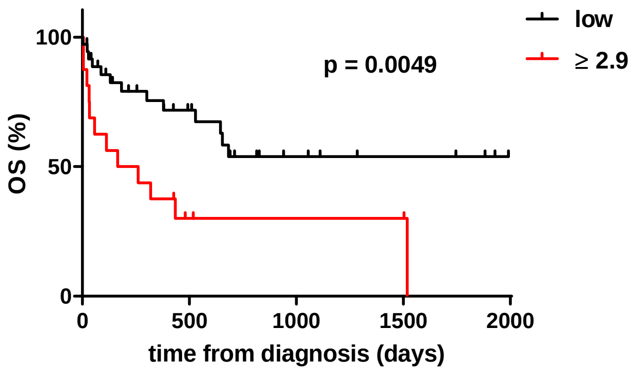


Figure S5: Impact of PD-L1 mRNA on patient survival

Survival analysis according to v1/v2 mRNA PD-L1.

A: overall survival (OS)

B: relapse-free survival (RFS).

A

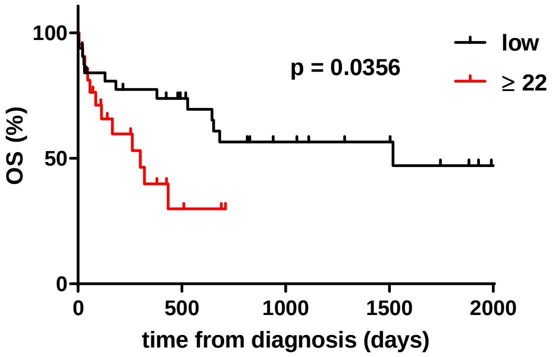


B

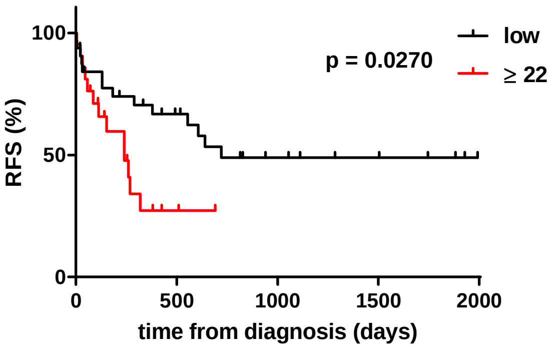
